# A reanalysis of Schaefer et al. does not indicate extensive CRISPR/Cas9 mediated off-target editing events

**DOI:** 10.1101/159608

**Authors:** Reynald M. Lescarbeau, Bradley Murray, Thomas M. Barnes, Nessan Bermingham

## Abstract

Recently Schaefer *et al.* (1) reported the presence of more than a thousand genomic differences between mice that had been edited with *S. pyogenes* Cas9 at the zygote stage, and a control mouse of the same strain. Given the overlap of genomic differences between the two edited mice that were not found in the control mouse, the authors concluded that these differences arose from a Cas9-dependent activity. We feel that this conclusion is inappropriate, for three key reasons: 1) there was incomplete analysis of the genetic variation; 2) there was no consideration that the variation is naturally arising in these animals; and 3) the inferred behavior of Cas9 lies outside of its understood mechanism of action. Attribution to Cas9 activity should require a burden of proof, that we believe has not been met.

---

The origin of the variation is a question of timing: the authors propose, given the pattern of homo- and heterozygosity, that it occurred within the first few divisions of the zygote as a result of the Cas9-mediated repair of the *rd1* allele of *Pde6b* (2), an ophthalmic disease target. The alternative explanation is that the genetic variability was not Cas9-mediated but rather was present in the FVB/NJ mouse lineage prior to the treatment of the animals. These two alternatives (Cas9 vs. background variation) can be distinguished by examining the pattern of shared and unique variants between the sequences of the mm10 reference genome (3), the FVB/NJ reference genome (4), the FVB/NJ control animal in Schaefer *et al.* (1), and the two edited mice in that study (F03 and F05).

We downloaded and reprocessed the Schaefer *et al.* data as well as the FVB/NJ whole genome sequencing data from the Wellcome Trust Sanger Institute (4) using an adaptation of the GATK best practices workflow (Supplemental Methods), leading to several conclusions. First, despite being extensively inbred, both the reference FVB/NJ genome (Fref), as well as the control FVB/NJ animal sequenced by Schaefer *et al.* (Fcon) have extensive heterozygosity (263,257 and 307,145 sites with differing SNVs from mm10 in Fref and Fcon, respectively, with 190,869 of these heterozygous sites overlapping between the two animals; Supplemental Table 1). Substantial allele imbalance in these variants was also noted across all three animals from Schaefer *et al.* raising concerns about the ability to call heterozygous variants accurately (Supp. Fig. 1). Nonetheless, this heterozygosity indicates that this line is far from “clonal”, which is not surprising given the expected inbreeding depression effects of genome-wide homozygosity. Second, based on publication timing between Wong *et al.* (4) and Schaefer *et al.* (1), several years elapsed between the birth of the Fref and Fcon animals, and during this time 640,780 SNVs differences appear between these two animals (459,980 homozygous and 180,800 heterozygous sites in either animal, using the other as reference). Third, the three mice in Schaefer *et al.* are clearly more related to each other than any is to Fref, sharing most of their differences from Fref. This is illustrated by the fact that of 96,243 sites which contain homozygous variants in Fcon where Fref was wild type, 91,975 of those are homozygous variants in F03 and 91,539 are homozygous variants in F05 (Fig. 1A and B). Fourth, while we don’t know whether Fcon was contemporaneous with F03 and F05, the genetic divergence of these 3 animals does not appear on its face to support the idea of a burst of induced variation by Cas9, especially given the amount of variation already present in the FVB/NJ line. While there may be a small excess of variations induced by Cas9 due to well-understood off-target mechanisms (5), the significant excess of background variation masks detection at the WGS level. No off-target or specificity analysis appears to have been performed on these guides prior to their use (2); what is required, and lacking, is the sequence of the parents of F03 and F05.

**Figure 1.**
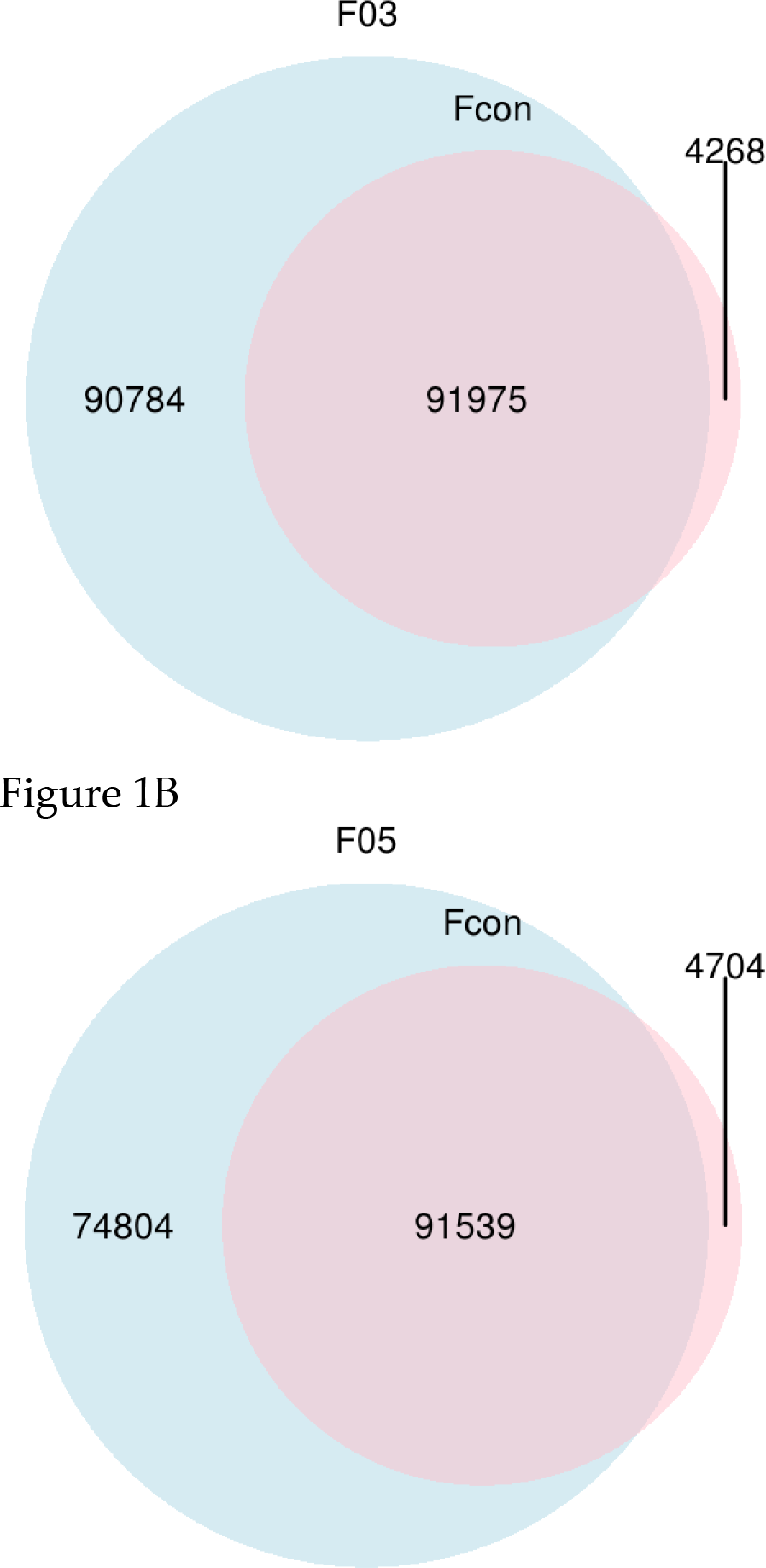
Homozygous variants found in edited mice and Fcon, which are wild type in Fref, are highly overlapping. A) The number of homozygous variants in F03 and the FVB/NJ control animal in Schaefer *et al.* (1) (Fcon) which are wild type in the reference FVB/NJ genome sequence (Fref) are depicted, as well as the number of overlapping variants in these sets. B) The number of homozygous variants in F05 and Fcon which are wild type in Fref are depicted, as well as the number of overlapping variants in these sets.

Notably, in an unrelated study, WGS of unedited inbred (except for a small region around *agouti*) C57BL/6J mouse littermates showed 985 sites of variation (SNVs and indels) between individuals (6), a number similar to that found in the authors’ dataset (696 SNVs and indels). Furthermore, a second unrelated study directly examined the effects of Cas9 editing (with intact Cas9 cleavase or Cas9D10A nickase) using WGS (7). These authors saw significant variation between individuals, but after comparison with relevant control parental and sib genomes, demonstrated the opposite result to Schaefer *et al.*, namely that Cas9 induced no unexpected mutations.

Beyond a lack of evidence of a recent origin of the variation, the authors’ explanation requires activities that Cas9 is not known to possess. First, the authors show that none of the edited sites has sufficient target homology to the guide to support Watson-Crick gRNA-DNA base pairing, a requirement for Cas9 activity (5). There was no genome-wide unbiased off-target analysis done with the chosen guide RNA (2), so we have no baseline for understanding its specificity. Second, 13/30 (43%) sites presented in Schaefer *et al.* lack a GG PAM within 4 bases of the variant, the PAM being the only known pre-requisite for Cas9 to engage with DNA (8). Third, Cas9 introduces indels, not SNVs, as a result of true off-target editing. There is no known mechanistic basis for wild type Cas9 to induce SNVs, nor did the authors propose one. Indeed, of the variation present, there is an excess of transitions over transversions, which is observed in naturally occurring variation (Supplemental Table 2) (9).

In summary, the principal conclusions of the correspondence (that the variation seen was unexpected, and that it was Cas9 mediated) are not supported by a reanalysis of their data. The unmanipulated FVB/NJ mouse line used in this study exhibits extensive heterozygosity, contrary to the authors’ assumption that the line is predominantly homozygous at all sites. Given this inherent variation, the expected finding would be that most variants are shared, but some end up only in certain individuals (6, 10), and that is what was seen.

## Data availability

The code used to analyze the data, along with the parameters used for variant calling, are available at github.com/Intellia/SchaeferAnalysis.

## Supplemental Methods

Sequencing reads were downloaded and split into pairs using the sra-tools toolkit and the following SRA accession numbers (SRR5450998, SRR5450997, and SRR5450996). Reads were aligned to the GRCm38/mm10 reference genome using the BWA-mem algorithm (v0.7). PCR and optical duplicates were the marked using Picard MarkDuplicates (v1.83). Base Quality Recalibration was done using the GATK BaseRecalibrator and known sites from mouse dbSNPv138 and the Mouse Genome Project (v3.4). Recalibrated BAM files the underwent variant calling using the GATK HaplotypeCaller. The FVN/NJ data was downloaded from the Wellcome Trust Sanger Institute Mouse Genomes Project and processed through the same pipeline. Sites were filtered for a minimum depth of 23, minimum quality of 30, and minimum genotype quality of 20. No other filtering was performed. At each stage of the reanalysis, select variants were manually inspected in a genome browser to confirm variants calls. Transition and transversion rates were calculated using vcftools on the VCFs provided by Schaefer et al (personal communication).

**Supplemental Figure 1.**
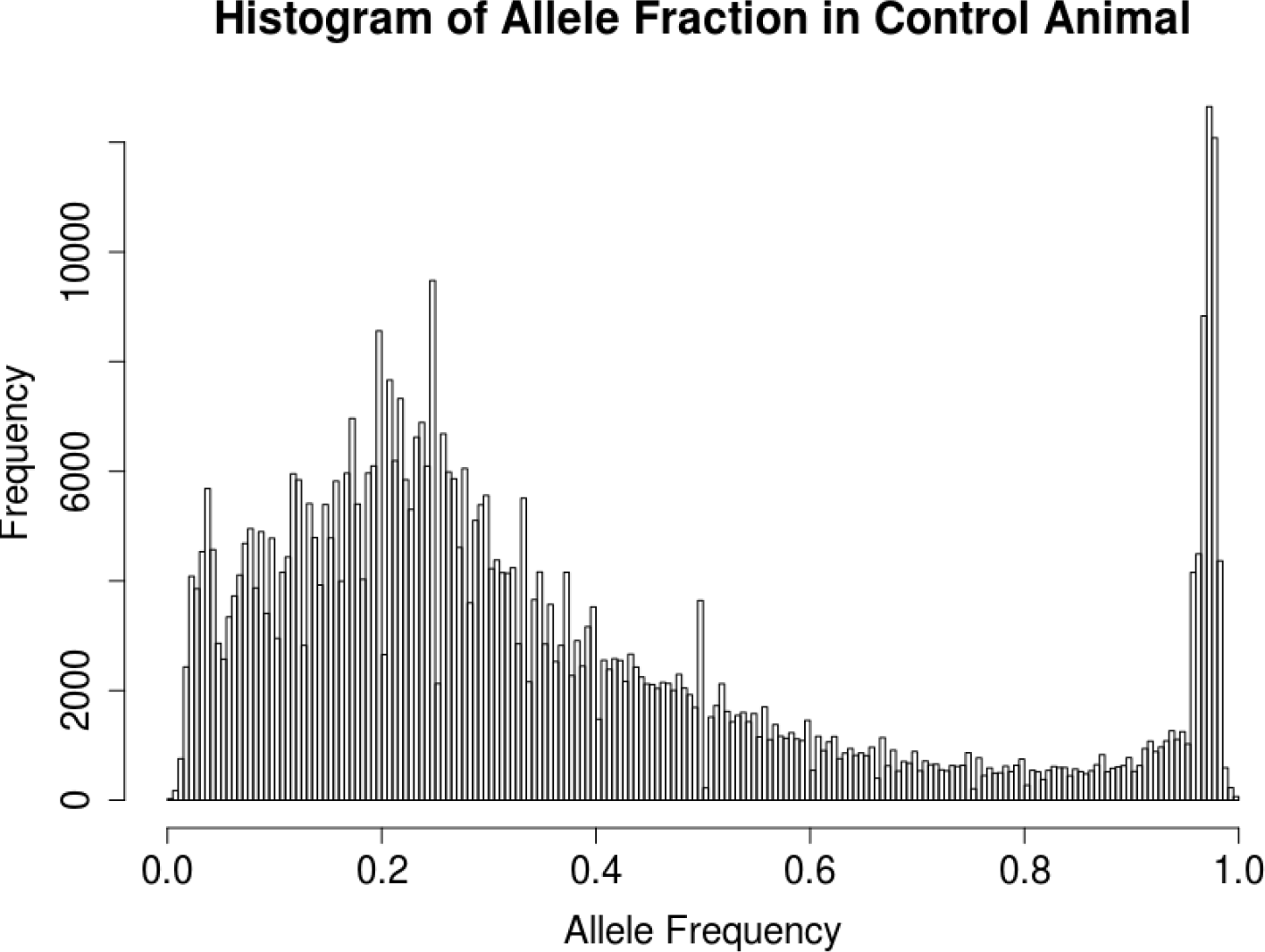
Frequency histogram of a randomly subsampled set of heterozygous variants from Fcon illustrating allele balance.

**Supplemental Table 1.** Twenty representative heterozygous SNVs present in the FVB/NJ reference animal and in the control animal sequenced by Schaefer *et al.* (1)

| FVB/NJ Reference Animal | Schaefer Control Animal |
| --- | --- |
| Chr7:11370951 | ChrX:170674818 |
| Chr17:38558017 | Chr4:69976351 |
| Chr11:71165478 | Chr12:114784612 |
| Chr6:129976386 | Chr2:177859988 |
| Chr11:91922049 | Chr12:114674982 |
| Chr7:11067972 | Chr10:22184941 |
| ChrX:85701552 | Chr11:27283274 |
| Chr12:113687320 | ChrX:75687910 |
| Chr7:103812725 | Chr4:60112555 |
| Chr17:36391328 | Chr12:18223142 |
| Chr17:48162392 | Chr1:84989932 |
| Chr12:40036401 | Chr6:130051235 |
| Chr6:130066283 | Chr17:38485149 |
| Chr7:42452171 | Chr6:128622254 |
| Chr4:20898259 | Chr1:177819208 |
| Chr5:15496745 | Chr14:4290977 |
| ChrY:10060668 | Chr12:115782701 |
| Chr12:22936103 | Chr6:130227066 |
| Chr6:130388137 | Chr13:113304009 |
| Chr14:43934645 | Chr4:147792139 |

**Supplemental Table 2.** Transition/Transversion ratios calculated on the VCFs provided by Schaefer *et al*.

|  | F03 vs Control | F05 vs Control |
| --- | --- | --- |
| Strelka | 1.31 | 1.28 |
| Mutect | 1.34 | 1.32 |
| Lofreq | 1.22 | 1.16 |
| Mean | 1.29 | 1.25 |

## References

1 Schaefer, K. et al. Unexpected mutations after CRISPR-Cas9 editing in vivo. Nat. Methods 14, 548–549 (2017).

2 Wu, W.-H. et al. CRISPR repair reveals causative mutation in a preclinical model of retinitis pigmentosa. Mol. Ther. 24, 1388–1394 (2016).

3 Mouse Genome Sequencing Consortium et al. Initial sequencing and comparative analysis of the mouse genome. Nature 420, 520–62 (2002).

4 Wong, K. et al. Sequencing and characterization of the FVB/NJ mouse genome. Genome Biol. 13, R72 (2012).

5 Doench, J.G. et al. Optimized sgRNA design to maximize activity and minimize off-target effects of CRISPR-Cas9. Nat. Biotechnol. 34, 184–191 (2016)

6 Oey, H. et al. Genetic and epigenetic variation among inbred mouse littermates: identification of inter-individual differentially methylated regions. Epigenetics & Chromatin 8, 54–65 (2015)

7 Iyer, V. et al. Off-target mutations are rare in Cas9-modified mice. Nat. Methods 12, 479 (2015).

8 Anders, C. et al. Structural basis of PAM-dependent target DNA recognition by the Cas9 endonuclease. Nature 513, 569–573 (2014).

9 Tubbs, A. and Nussenzweig, A. Endogenous DNA damage as a source of genomic instability in cancer. Cell 168, 644–656 (2017).

10 Shorter, J.R. et al. Male infertility is responsible for nearly half of the extinction observed in the mouse collaborative cross. Genetics 206, 557–572 (2017).

## Supplemental References

1. McKenna, A. et al. The Genome Analysis Toolkit: a MapReduce framework for analyzing next-generation DNA sequencing data. Genome Research 20, 1297–303 (2010).

2. Danecek, P. et al. The variant call format and VCFtools. Bioinformatics 27, 2156–2158 (2011).

3. Li, H. and Durbin, R. (2009) Fast and accurate short read alignment with Burrows-Wheeler Transform. Bioinformatics, 25:1754–60

